# SAXSDom: Modeling multi-domain protein structures using small-angle X-ray scattering data

**DOI:** 10.1101/559617

**Authors:** Jie Hou, Badri Adhikari, John J. Tanner, Jianlin Cheng

## Abstract

Many proteins are composed of several domains that pack together into a complex tertiary structure. Some multidomain proteins can be challenging for protein structure modeling, particularly those for which templates can be found for the domains but not for the entire sequence. In such cases, homology modeling can generate high quality models of the domains but not for the assembled protein. Small-angle X-ray scattering (SAXS) reports on the solution structural properties of proteins and has the potential for guiding homology modeling of multidomain proteins. In this work, we describe a novel multi-domain protein assembly modeling method, SAXSDom, that integrates experimental knowledge from SAXS profiles with probabilistic Input-Output Hidden Markov model (IOHMM). Four scoring functions to account for the energetic contribution of SAXS restraints for domain assembly were developed and tested. The method was evaluated on multi-domain proteins from two public datasets. Based on the results, the accuracy of domain assembly was improved for 40 out of 46 CASP multi-domain proteins in terms of RMSD and TM-score when SAXS information was used. Our method also achieved higher accuracy for at least 45 out of 73 multi-domain proteins according to RMSD and TM-score metrics in the AIDA dataset. The results demonstrate that SAXS data can provide useful information to improve the accuracy of domain-domain assembly. The source code and tool packages are available at http://github.com/multicom-toolbox/SAXSDom.

## 1. Introduction

Protein domains, as fundamental units of protein structure, have been extensively studied and commonly identified in most proteins, especially in large eukaryote proteins ^1^. During the process of evolution, domains are independently folded with particular functions, and arranged with different combinations to form various new proteins ^2^ Domains in a protein are generally connected by intrinsically flexible and unstructured linkers ^3^ while sometimes one domain is inserted into intra of another domain ^4^. Domain linkers have been shown as an essential role in maintaining domain-domain interaction, protein stability and domain-domain orientation ^5–6^. Understanding the domain arrangement is central to interpreting the structure, function and evolution of the multi-domain proteins ^7^. Thus, domain identification is generally an important first step in computational protein three-dimensional structure prediction ^8–9^. With the wide application of experimental techniques such as X-ray crystallography and NMR spectroscopy, the number of determined protein structures is exponentially growing and provides a valuable resource for bioinformatics research ^10^. Consequently, the computational protein structure prediction from sequences has advanced greatly in the last decades ^9, 11–17^. Most computational methods start with parsing sequences into domains and perform either comparative (homology) structure modeling ^9, 18^ or *de novo* structure prediction ^8, 19^ on individual domains. The predicted structures of domains are subsequently assembled into full-length structural model in terms of free energies ^11, 20^. In the *de novo* structure prediction, different free energy functions are frequently taken into account when the optimization approaches are applied to fold a protein structure into native-like state that has lowest energy ^11, 19–21^. Several potential functions have been widely considered, such as Van der Waals, hydrophobic effects, electrostatic interactions, hydrogen bonding propensity, and backbone interaction potential ^11^. The changes of free energy during the conformation sampling were generally captured by different optimization approaches, such as Monte Carlo simulation ^11^, simulated annealing ^19^ and gradient descent ^20^. The integration of energy functions with optimization techniques facilitates yielding native-like conformation. Despite showing the effectiveness, it remains to be challenging and difficult in sampling large conformational space for *ab initio* protein structure prediction, especially for those proteins with large protein size, or presence of multi-domains for which templates can be found for the domains but not for the entire sequence. Therefore, different kinds of experimental information were investigated to assist the computational modeling by introducing additional energies ^22–24^.

Unlike the high-resolution structure determination, the small-angle X-ray scattering (SAXS) technique can detect low-resolution structural information (i.e. 12-20 Angstrom) in solution without crystal formation and can be efficiently and rapidly determined with lower costs ^25–26^. Many advances have been made in the use of SAXS information for structural analysis in recent years, including processing and analyzing SAXS data ^27–29^, and the applications that integrate SAXS information with computational approaches in protein structure modeling ^30–33^. Compared with the accurate determination of atomic coordinates using X-ray or NMR, SAXS provides the statistics of all the electron pair distance for a protein in solution. Consequently, we can derive distance distribution of all pair atoms, mass size and shape of proteins from SAXS data. Thus, it is promising to use SAXS data in the protein structure modeling as a prior experimental restraint, especially useful for the study of domain-domain interaction and protein-protein docking ^23^. In the recent Critical Assessment of Protein Structure Prediction (CASP) competition, SAXS information was incorporated into the data-assisted category that aimed to assess the potential of integrating SAXS data with protein structure prediction methods for protein folding ^24^. Most CASP12 approaches utilized SAXS as additional driving restraints, which include (1) *χ* score that measures the fitness of scattering curves between the experimental SAXS intensities and computed intensities from the model; (2) pair distance distribution (*P*(*r*)) that quantitates the histogram matching of atom pair distance; (3) Radius of Gyration (*Rg*) that restraints the structural size. Even though the SAXS data holds great promise for accurate structural folding, it is still challenging to leverage the different types of experimental knowledge in the application of computational modeling.

In this work, we investigated whether incorporating the restraints from SAXS data can improve the multi-domain assembly. We developed a novel framework that systematically integrates the probabilistic approach with SAXS data to assist in structure folding and multi-domain assembly. Our method applies probabilistic Input-Output Hidden Markov model and Monte Carlo sampling to simulate the domain-domain orientation with enforced energies derived from SAXS data, so that it can generate near-native structures that have low free energy and close SAXS profile match. In addition, we examined the correlation between the several SAXS scoring functions and structural qualities (i.e. RMSD) on the CASP proteins, which shows the effectiveness of SAXS data in the structural analysis. Our method shows a significant improvement in domain assembly and structure folding after incorporating SAXS information as additional energies to the physical-based force field, and demonstrate its promising use of SAXS data in computational protein structure modeling.

## 2. Method

### 2.1. Benchmark sets

To assess how well each SAXS energy correlates with structural qualities (i.e. RMSD) ^34^, we collected predicted structural models generated for protein targets that were tested in the 8^th^, 9^th^, 10^th^, 11^th^ Critical Assessments of Structure Prediction (CASP) experiments ^35^. The proteins whose experimental structures were available were selected for preliminary analysis. The dataset contains 112,050 models corresponding to 428 single-domain and multi-domain proteins. The detailed statistics for this dataset is provided in **Table S1**.

In addition, we evaluated our proposed method on the two types of datasets to validate the effectiveness of SAXS data in protein multi-domain assembly. The first dataset contains multidomain proteins from CASP8-12 whose experimental structures are available to date. The domain definition (i.e. number of domains and the domain boundaries) of each protein was determined by CASP assessors ^36^. Since our method requires the continuous domains as input, the domains with chain-break (i.e. distance of adjacent Ca-Ca atoms is larger than 4 Å) were removed from the dataset. Finally, we collected 51 CASP multi-domain proteins for the domain assembly analysis. The length of domain linkers among 51 proteins ranges from 5 to 21. We randomly selected 5 targets to determine the weights for the SAXS energy terms in the energy function. The remaining 46 targets were used to compare the performance of different SAXS scoring functions for domain assembly. The structures of individual domains for all 51 CASP targets were directly derived from their native protein structures and were further used in domain assembly.

The second dataset is a collection of two-domain proteins curated in the AIDA domain assembly method ^21^. The number of domains in each protein was determined by DomainParser ^37^. Unlike using the native domain structures for assembly in the CASP dataset, we first used MULTICOM tertiary structure system ^38^ to predict the structures of individual domains of proteins from their homology templates. The domains whose predicted structures have TM-score > 0.9 against their native structures were selected for domain assembly. Finally, MULTICOM successfully predicted high-quality models for domains of 73 proteins in the AIDA dataset. The length of domain linkers in 73 proteins ranges from 5 to 15. The predicted structures were used for domain assembly analysis.

### 2.2. Domain-Domain orientation driven by united-residue model and probabilistic sampling

Given individual domain structures for a protein sequence, our method first converts the polypeptide chains of domains into united-residue representation as described in the UNRES model ^19, 39^. In the united-residue model, the backbone of the polypeptide chain is constituted by a sequence of alpha-carbon atoms linked by virtual bonds, and the conformation of the protein chain is determined by virtual-bond length (*b_ca_i__*), virtual-bond angles (*θ_i_*), virtual bond dihedral angles (*τ_i_*) among adjacent backbone alpha-carbon atoms. In addition, the united side-chains are attached to the alpha-carbon atoms where two side-chain angles (*δ_i_* and *γ_i_*) and virtual-bond length (*b_SC_i__*) determine the location of side-chain with respect to the alpha-carbon atoms in the backbone. The six variables parameterize the geometry of alpha-carbon (*Cα_i_*) and side-chain (*SC_i_*) at i^th^ residue of a polypeptide chain in the conformation space. We used Input-Output Hidden Markov Model (IOHMM) that was pre-trained in our previous work ^19^ to sample the virtual-bond lengths and virtual-bond torsion angles given amino acids and predicted secondary structure in the linker regions. Each cycle of Monte Carlo sampling generates one acceptant movement for domain-domain orientation using simulated annealing algorithm. It is worth noting that the structural conformation of individual domains keeps unchanged during sampling, so that the conformation of linker regions can be conditionally resampled given the known prior structural information of domains based on the probabilistic model, which can predict more accurate local structural preferences of linkers than random sampling and potentially reduce the number of local movements in the conformational space.

Our method implements the domain assembly based on the following steps. Given the full-length sequence of a protein, our method first predicts its 8-class secondary structure using SSpro ^40^, and samples the united-residue conformation for the entire polypeptide chain using IOHMM model for structure initialization. After the conformation is initialized, the torsion angles and virtual-bond lengths of alpha-carbon and its side-chain atoms at each position of residues in the full-length polypeptide chain are updated according to their geometry in the predetermined domain structures. The regions whose structure information are not provided in the provided domain structures are considered as linkers that anchors domains together. The conformation of linker regions is then sampled using IOHMM model and orients the domain structures using simulated annealing algorithm to generated structure models with lowest structural energy, as depicted in the **Figure 1**. Therefore, our method can be applied to assemble any number of domains for multi-domain proteins.

**Figure 1.**
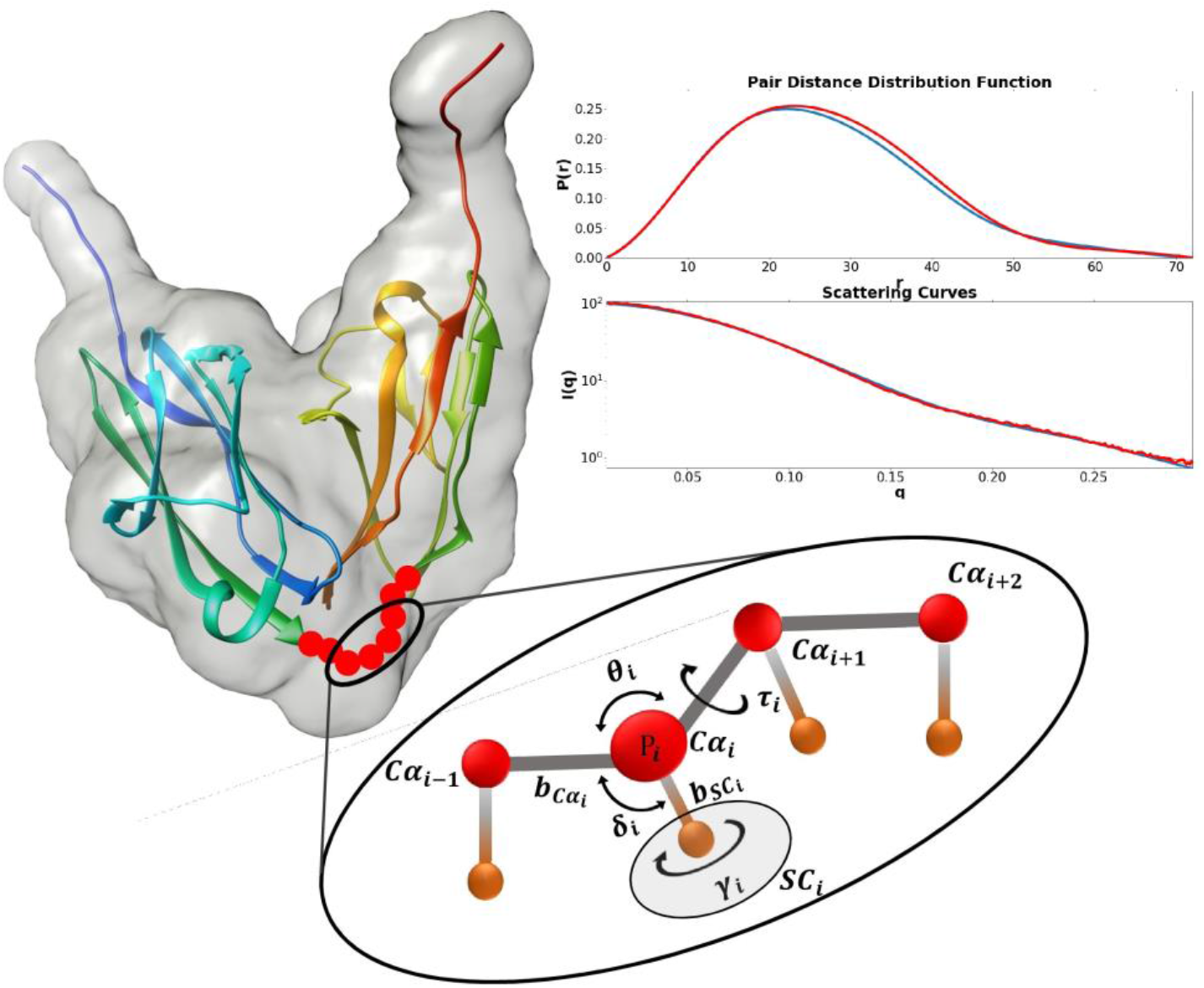
Parameterization of conformation in linker regions and overall shape match with SAXS data.

### 2.3. Integrating physics-based force field with SAXS restraints for domain-domain assembly

Our method adopts the same energy configuration using united-residue physics-based force field that was defined in our previous work ^19^ to represent the energy of a united-residue peptide chain. The physics energy includes the mean free energy of hydrophobic (hydrophilic) interactions between side chains (*E_SC_i_SC_j__*), excluded-volume potential of side-chain and peptide group interaction (*E_SC_i_p_j__*) and the backbone peptide group interaction to represent the average electrostatic interactions(*E_p_i_p_j__*) for any pair residues in the *i^th^* and *j^th^* positions in the polypeptide chain, as represented in Equation (1)

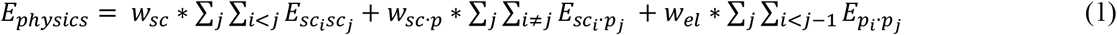

Unlike our earlier approach that generated chain conformation based on stepwise sampling of foldon units, our method only samples the conformation of the linker regions and keeps the geometry of domain structures fixed. Therefore, the physics-based force field of intra-domain interactions is stable during conformation sampling, and the energy of chain conformation is only affected by the interactions of all inter-domain residues (i.e. interaction interface) and all linker residues, where the physics energy can be further represented as:

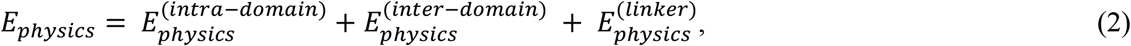

It is worth noting that the energy of hydrophobic (hydrophilic) interactions between side chains of linker residues plays an important role in the protein folding and domain-domain movement ^41^. Studies showed that the average residue hydrophobicity (hydrophilicity) is largely influenced by the size of linkers, where longer linkers are more hydrophilic and exposed so that they induced larger domain motions in the conformation space. Inversely, smaller linkers showed more hydrophobic which may significantly restrain the domain-domain movement ^6^.

In order to assist the domain-domain orientation using knowledge-based information, we introduced additional energy terms corresponding to the SAXS restraints for the total energy calculation, defined as:

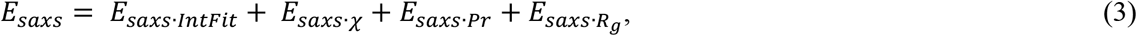

where *E_saxs·IntFit_* represents the normalized fitness between the experimental SAXS intensity and computed intensity from the models, which is defined as:

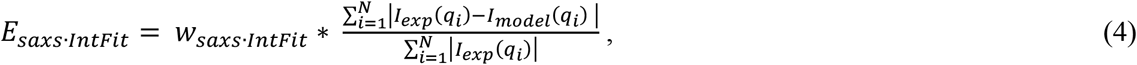

where *I_exp_*(*q*) and *I_model_*(*q*) correspond to the experimental SAXS intensities derived from SAXS profile and estimated intensities from decoys, respectively. We employ the same strategy as FoXS ^28, 42^ proposed to calculate best fitted intensity profile *I_model_*(*q*) by minimizing the *χ* function as follows:

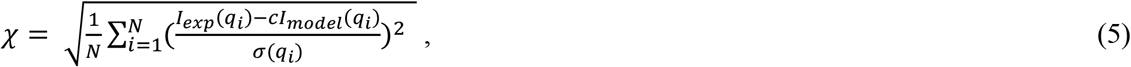

where *σ*(*q*) is the experimental error of the measured SAXS profile. *N* is the number of points in the SAXS profile, and *c* is the scale factor which is determined by performing the linear least-squares minimization to derive minimum value of *χ*. Therefore, our method also included *χ* as additional score term to account for the energic contribution of SAXS profile matching degree, which is defined as:

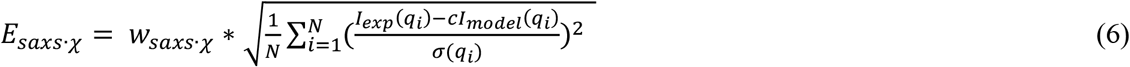

*E_saxs·Pr_* represents the Kullback-Leibler divergence between the pairwise atom-atom distance distribution function *P*(*r*) derived from SAXS profile and the pair distance distribution computed from the model, which is defined as:

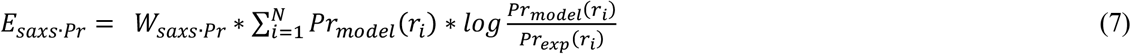

Compared to intensity profiles, *P*(*r*) provides more intuitive information about the shape and size molecules. It is straightforward to derive maximum distance (*d*_max_) and estimated *P*(*r*) based on the inverse Fourier transformation of Debye formula given SAXS intensity profiles ^29, 43^. The pair distance distribution of the protein structure is directly calculated from its atomic coordinates. The last energy term *E_saxs·Rg_* represents the radius of gyration restraints, defined as

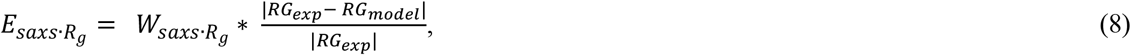

which measures the agreement between radius of gyrations estimated from SAXS data and calculated radius of gyration from protein model. The SAXS-related information (i.e. SAXS intensity, pair distance distribution and radius of gyration) described above was derived via algorithms as implemented in Integrated Modeling Platform (IMP) package ^44^ for SAXS energy calculation.

We adopted the same weight configuration for physics-based force field energy terms listed in Eq (1) as our previous method ^19^, where *w_sc_* = 1.00000, *w_sc·p_* = 2.73684, and *w_el_* = 0.06833. For the SAXS energy terms described in the Eq (3), we set *w_χ_* = 10.0, *w_saxs·fit_* = 700.0, *W_saxs·Pr_* = 700.0, and *w_saxs·Rg_* = 700.0 after experimenting with several weights on the small training proteins.

Therefore, the energy for the multi-domain polypeptide chain used in our method is defined as:

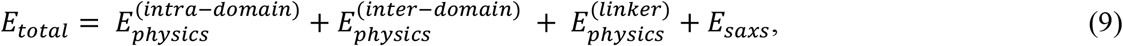

In addition to the four SAXS-related scoring functions as defined in Equation (4,6,7,8), we also implemented additional 10 SAXS-based score functions to account for the matching degree between experimental SAXS profiles and computed profiles from models, as shown in **Table S1**.

Since the physics-based energies are calculated from the united-residue models, however, the SAXS energies calculation requires the full-atom representation with at least Ca-trace included. We reconstruct the Ca-trace and side-chain from the united-reside protein representation by using PULCHRA ^45^ to generate full-atom protein models for SAXS energy calculation. In order to speed up SAXS fitting and computation, the functions of FoXS ^42^, PULCHRA ^45^ and IMP ^44^ have been built in our system instead of calling them as external programs during sampling.

We used the Monte Carlo simulation with simulated annealing algorithm for energy minimization to search the lowest-energy assembled multi-domain conformation. Since only linker regions are resampled during domain-domain orientation, the sampling space is significantly reduced. The number of Monte Carlo cycles for each linker is set to the number of residues in linker times 100. Given an assembled protein decoy in each cycle, the total energy including physical interaction energies and SAXS fitting energies is calculated and compared to the energy of previous conformation. The domain movement is accepted or rejected according to the probability proportional to 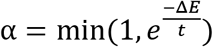, where the Δ*E* represents the energy change for each domain movement, and t is the temperature of simulated annealing.

## 3. Results

### 3.1. Evaluation of different SAXS profile matching score functions

In this study, we first examined the correlation between different score functions derived from SAXS profile with the structural quality of a given protein model. Total 14 score functions were considered to account for the SAXS profile matching degree between the computed intensities derived from the model and SAXS intensities from the experimental data, as shown in **Table S2**. Since Root Mean Square Deviation (RMSD) aims to measure the distance variation between Cα atoms of superimposed protein structures and has been commonly used to validate if the multiple domains have been correctly assembled (e.g. RMSD < *3 Å*) ^21, 46^, we evaluated the correlation (i.e. Pearson correlation coefficient) between the 14 SAXS-based scoring functions and RMSD metric on the predicted server models of 428 proteins from CASP8 to CASP11. Given a predicted model and its native structure, the profile I(q) was derived using FoXS program ^42^, and pair distance distribution function P(r) was obtained using inverse Fourier transformation from the intensity profile I(q) by GNOM ^47^. The intensity profile estimated from the experimentally determined structure was considered as theoretical profile for the protein. For each predicted model, we generated SAXS intensity profiles I(q) from both full-atom and Cα-atom protein structure and then calculated the SAXS matching scores according to the 14 score functions as defined in **Table S2**. The structural quality of each predicted model was evaluated according to RMSD metric by superimposing the predicted structure with its native structure. The Pearson correlation coefficient (PCC) between the RMSD and each of 14 SAXS scores of all predicted models for each protein was calculated and the averaged correlation of 428 proteins was summarized in the **Table S2** and **Table S3** in terms of full-atom and Cα-atom SAXS profile. As shown in the table, the divergence of pair distance distribution function P(r) (score 2), agreement of radius of gyration (score 3), and normalized fitness between computed and experimental SAXS intensities (score 5) showed the highest correlation with RMSD, with 0.6, 0.7, 0.59 for full-atom SAXS profile and 0.58, 0.66, 0.52 for Cα-atom SAXS profile respectively. The analysis demonstrates that the three score functions derived from Cα-atom SAXS profile doesn’t affect the correlation largely compared to the full-atom SAXS profile. This is also useful for Cα-trace backbone modeling because full-atom reconstruction is not required to generate SAXS profile to measure the fitness of computed and experimental intensities during the simulation, which facilitates the speed of model sampling and exploration of larger conformation space. Since χ function is the most common metric for comparison of scattering curves for SAXS, we also included it for analysis even though χ score (score 1) achieved relatively low correlation with 0.47 for full-atom and 0.38 for Cα-atom SAXS profile as shown in **Table S2** and **S3**. Therefore, in this work, we considered four score functions as SAXS energies as defined in Equation (4, 6, 7, 8). The correlation between the four SAXS scores and RMSD on CASP dataset is visualized in **Figure 2**.

**Figure 2.**
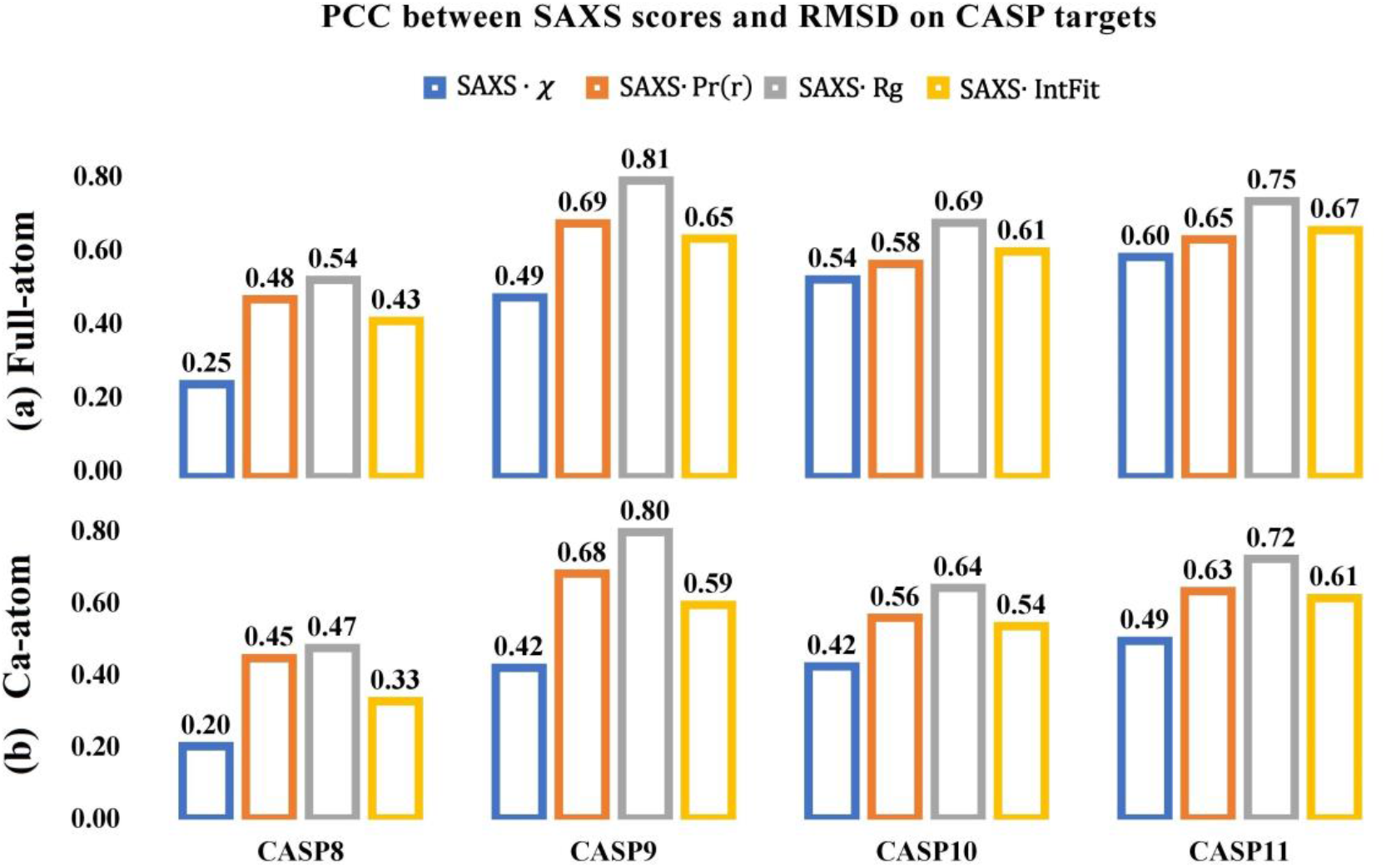
Pearson correlation coefficient of the structural quality (RMSD) against the SAXS score functions derived from (a) Full-atom and (b) Ca Atom of protein structure. Analysis was done based on the predicted models from CASP8-11.

### 3.2. Performance of SAXSDom in CASP multi-domain proteins using native domain structures

In order to validate the improvement of domain assembly after incorporating the information of SAXS data, we first developed a baseline approach, SAXSDom-abinitio, which used the basic united-residue physical based force field energies as defined in the Equation 1. We then developed 5 approaches that adopted different SAXS energy terms as defined in Equation (3, 4, 6, 7, 8), which were labeled as SAXSDom(*E_saxs·χ_*), SAXSDom(*E_saxs·IntFit_*), SAXSDom(*E_saxs·Pr_*), SAXSDom(*E_saxs·Rg_*) and SAXSDom(*E_saxs_*). All SAXSDom methods were employed to assemble domains for 46 CASP multi-domain proteins and each method generated 50 full-length decoys for each protein. For each protein, the initial structure of each domain was directly derived from its native structure, and the secondary structure of full-length protein sequence was predicted by SCRATCH ^48^. The theoretical SAXS intensity profile was calculated by FoXS from its native structure. After 50 decoys were generated, we used model quality assessment method Qprob ^49^ to rank the assembled models. Each method was evaluated based on the averaged TM-score and RMSD of top one, best of top five, best of top 50 models for all 46 proteins. The results for six methods are reported in the **Table 1**. When top one, best of top five and best of 50 models were evaluated, the averaged TM-score and RMSD of assembled models which were generated by different SAXS scoring functions were significantly improved compared to the performance of *ab initio* approach SAXSDom-abinitio. The P-value for the difference between the SAXS-based method and *ab initio* modeling according to TM-score and RMSD was reported in **Table S4**. For instance, the method SAXSDom(E_saxs_), which combines all four SAXS energy terms during conformation sampling, outperforms the method SAXSDom-abinitio by 9.97%, 11.98%, 11.37% of TM-score and 38.54%, 46.18%, 46.74% of RMSD for top one, best of five, and best of 50 models respectively. **Figure 3** shows the performance of five SAXSDom methods with different SAXS energies and SAXSDom-abinitio method evaluated on the best of 50 assembled models based on the RMSD, TM-score, and SAXS *χ* score. According to the evaluation, as shown in **Figure 3(A)**, the method SAXSDom(*E_saxs_*) outperforms the SAXSDom-abinitio in 40 out of 46 proteins in terms of RMSD and TM-score. We also evaluated the distribution of SAXS *χ* scores for all generated models. As expected, the SAXS *χ* scores of assembled models using SAXS information were lower than that of models built by *ab initio* sampling. As shown in the plot, the distribution of SAXSDom(*E_saxs_*) consistently shifted to lower SAXS *χ* score compared with SAXSDom-abinitio. **Figure 3 (B), (C), (D) and (E)** show the performance of domain assembly using four individual SAXS energy terms and their comparison with performance of ab initio sampling. The results clearly show that incorporating SAXS information as additional energies for conformational sampling can improve the accuracy of the domain assembly and protein folding. The results of method comparison evaluated on the top one and best of five assembled models are shown in **Figure S1** and **S2**.

**Table 1.** Summary of the domain assembly performance using *ab initio* modeling and different SAXS-related score functions on the 46 proteins in CASP dataset

| ScoreFunction | Top one |  | Best-of-Five |  | Best-of-50 |  |
| --- | --- | --- | --- | --- | --- | --- |
|  | TM-score | RMSD | TM-score | RMSD | TM-score | RMSD |
| SAXSDom-abinitio | 0.7265 | 8.4127 | 0.7580 | 6.4692 | 0.7959 | 4.4317 |
| SAXSDom($E_{saxs\cdot\chi}$) | 0.8117 | 5.0931 | 0.8518 | 3.4906 | 0.8775 | 2.6021 |
| SAXSDom($E_{saxs\cdot IntFit}$) | 0.7568 | 6.7671 | 0.8246 | 3.9591 | 0.8652 | 2.7355 |
| SAXSDom($E_{saxs\cdot Pr}$) | 0.8022 | 5.2716 | 0.8480 | 3.4606 | 0.8857 | 2.2904 |
| SAXSDom($E_{saxs\cdot Rg}$) | 0.7727 | 6.1951 | 0.8123 | 4.1972 | 0.8457 | 3.0301 |
| SAXSDom($E_{saxs}$) | 0.7989 | 5.1704 | 0.8488 | 3.4815 | 0.8864 | 2.3605 |

**Figure 3.**
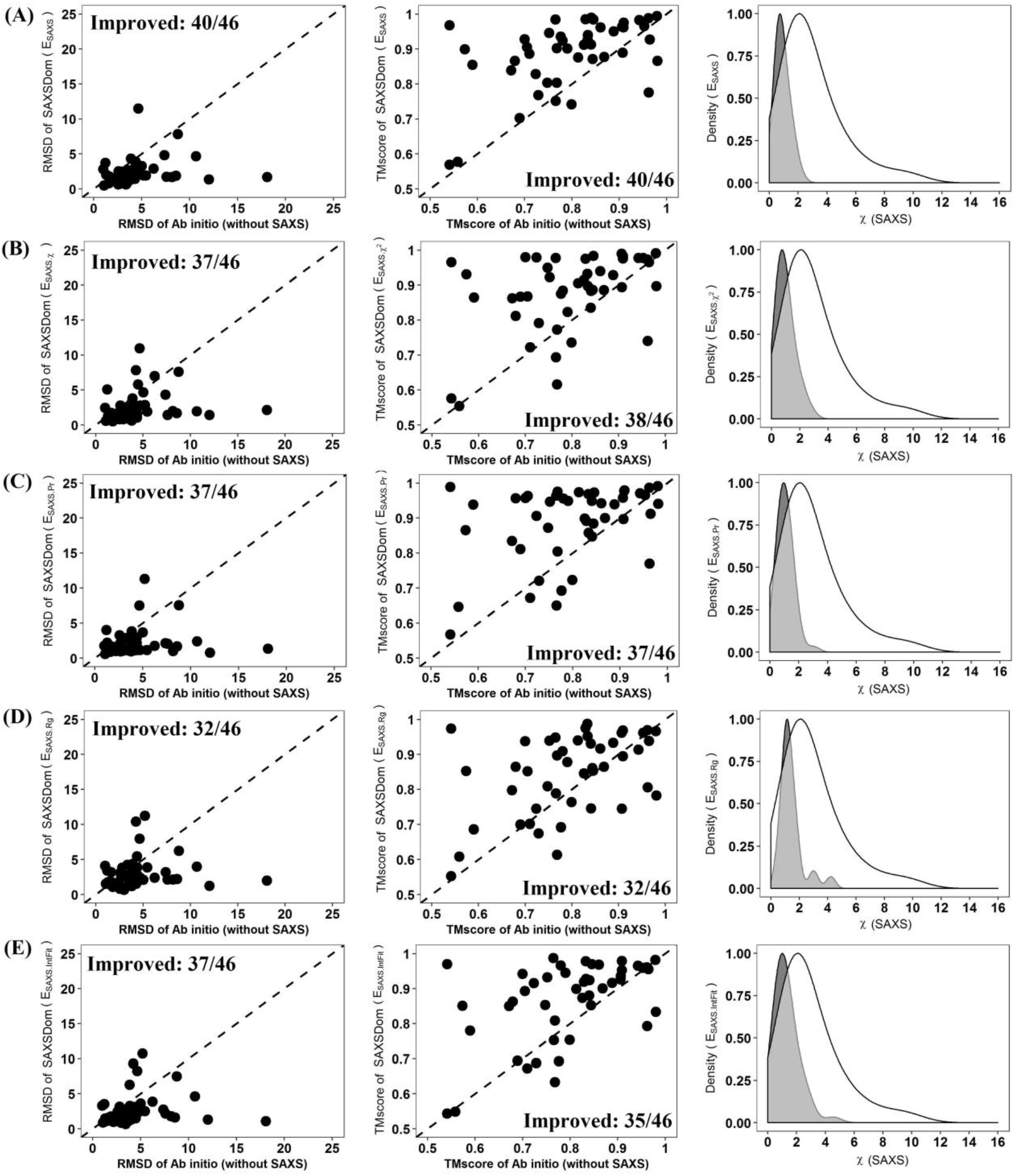
Comparison of five SAXSDom methods with SAXSDom-abinitio method on the best of 50 assembled models. **(A)** SAXSDom (*E_saxs_*) versus SAXSDom-abinitio (Left plot: TM_scores of SAXSDom (*E_saxs_*), models versus TM_scores of SAXSDom-abinitio models; Middle plot: RMSD of the models of the two methods; Right plot: Distribution of χ score of all assembled models for 46 proteins by two methods). **(B)** SAXSDom (*E_saxs·χ_*) versus SAXSDom-abinitio. **(C)** SAXSDom (*E_saxs·Pr_*) versus SAXSDom-abinitio. **(D)** SAXSDom (*E_saxs·Rg_*) versus SAXSDom-abinitio. **(E)** SAXSDom (*E_saxs·IntFit_*) versus SAXSDom-abinitio.

### 3.3 Performance of SAXSDom in AIDA multi-domain proteins using predicted domain structures

We also assessed the performance of SAXSDom using 73 multi-domain proteins which were originally curated by *ab initio* domain assembly approach AIDA ^21^. In our work, the domain structures for these 73 proteins were predicted by MULTICOM tertiary structure prediction method and then further assembled using our model protocol. SAXSDom then generated 50 assembled decoys using the reference SAXS intensities derived from the native structures of full-length proteins and applied Qprob to re-rank the models. The same protocol was applied to SAXSDom-abinitio to generate 50 decoys for the 73 proteins. The accuracy of top ranked models (i.e. top 1, best of five, best of 50 models) were subsequently evaluated according to TM-score and RMSD. We also compared our methods with another two state-of-art structure modeling approaches, Modeller ^20^ and AIDA ^21^. For each protein, Modeller and AIDA also generated 50 decoys which were ranked according to their default energies. The qualities of top ranked models generated by Modeller and AIDA were also evaluated and compared to our methods. **Table 2** reports the averaged TM-score and RMSD of top ranked models (i.e. top 1, best of five, best of 50 models) generated by four methods. As shown in the table, AIDA achieved relatively better performance in domain assembly compared to other three methods. The main difference between AIDA method and our approach is that AIDA represents the protein structure as backbone geometry defined by all backbone atoms and the energy potentials are derived from the main-chain atoms and side-chain, while SAXSDom adopted united-residue representation that describes the structures as a sequence of alpha-carbon (Ca) atoms with attached united side-chain and the energy is a basic implementation of united-residue physics based force field. However, SAXSDom still outperforms SAXSDom-abinitio and Modeller in terms of all metrics with statistically significance adopted by one-sample paired t-test, which also demonstrated the improvement by incorporating the SAXS restraints in domain assembly. **Figure 4** shows the performance of SAXSDom with SAXSDom-abinitio, AIDA and Modeller evaluated on the best of 50 assembled models based on the RMSD, TM-score, and SAXS *χ* scores. According to the evaluation, as shown in **Figure 4(A)**, the method SAXSDom outperforms the SAXSDom-abinitio in 50 out of 73 proteins in terms of RMSD and 45 out of 73 proteins in terms of TM-score. **Figure 4(B)** shows the performance of SAXSDom and AIDA that AIDA method was able to assemble domains with slightly better qualities according to RMSD, while, SAXSDom can generate assembled decoys that were better matched to the SAXS profile. **Figure 4(C)** shows that SAXSDom can generate significantly better models with lower SAXS *χ* scores compared to that of Modeller. The results of method comparison evaluated on the top one and best of five assembled models were also shown in **Figure S3** and **S4**.

**Table 2.**
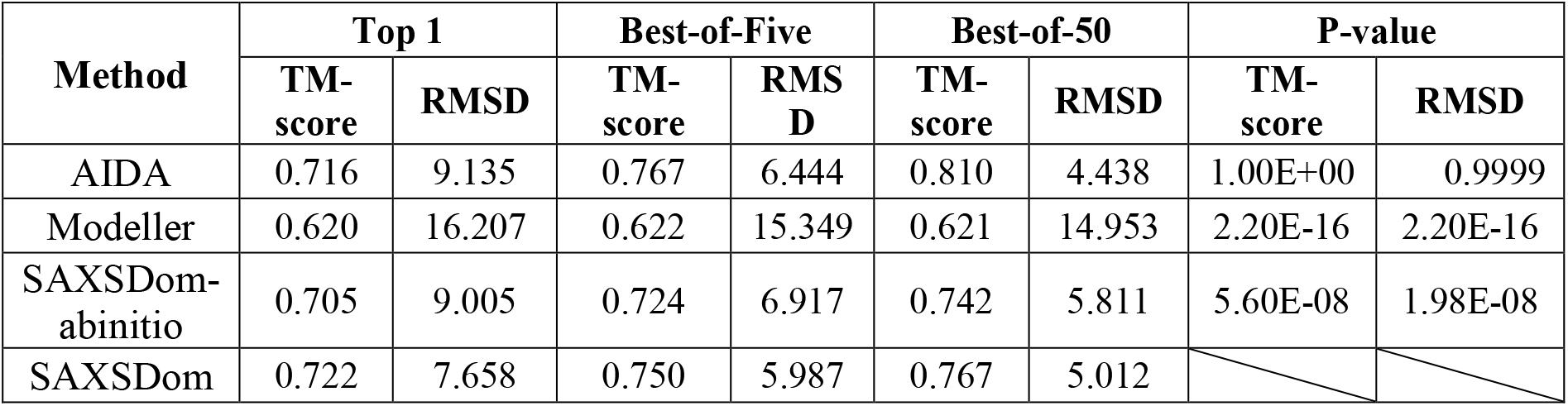
Summary of the domain assembly performance using for domain assembly methods on the 73 proteins in AIDA dataset

**Figure 4.**
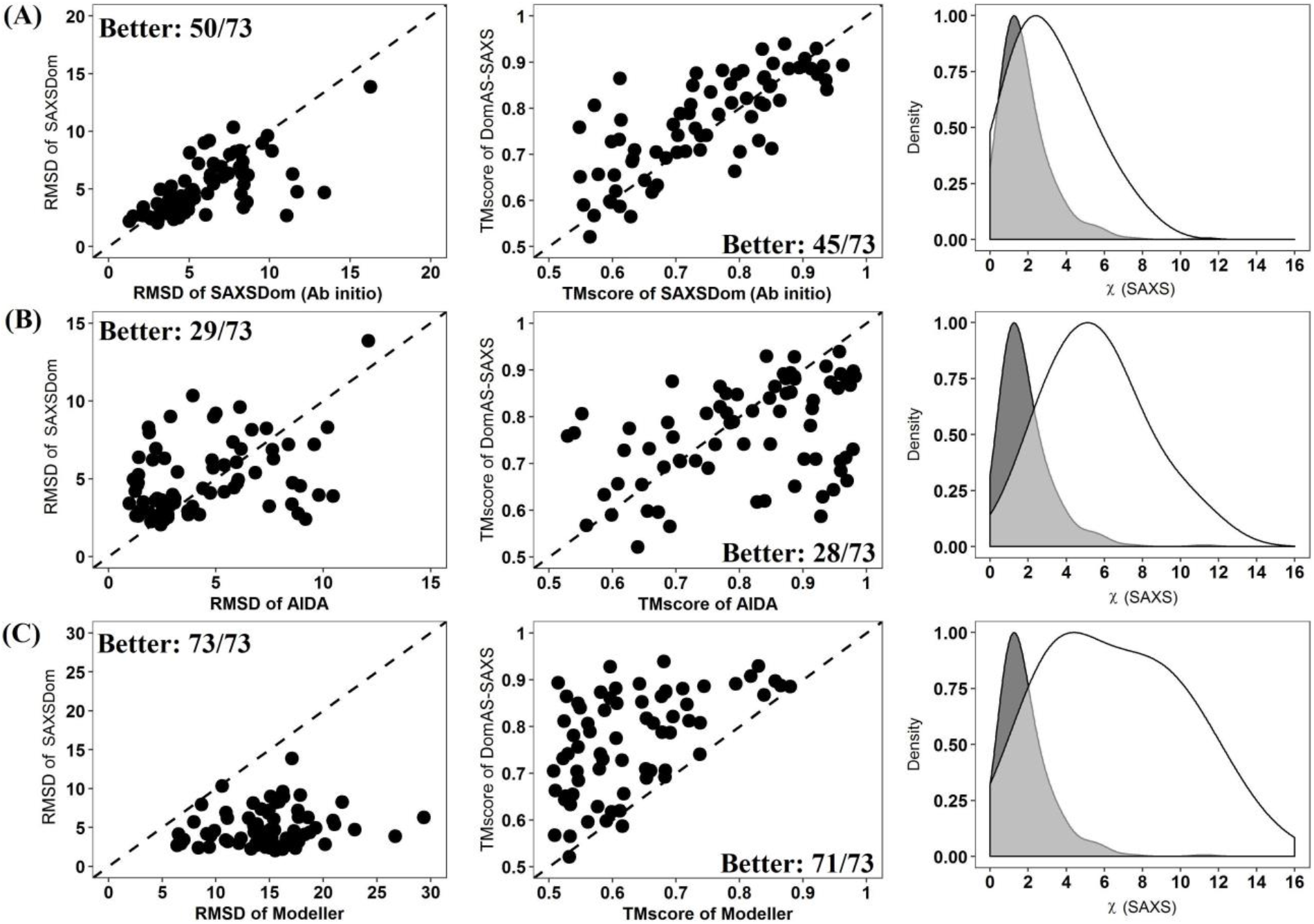
Comparison of SAXSDom with SAXSDom-abinitio, AIDA and Modeller on the best of 50 assembled model. **(A)** SAXSDom versus SAXSDom-abinitio (Left plot: TM_scores of SAXSDom models versus TM_scores of SAXSDom-abinitio models; Middle plot: RMSD of the models of the two methods; Right plot: Distribution of *χ* scores of all assembled models for 46 proteins by two methods). **(B)** SAXSDom versus AIDA. **(C)** SAXSDom versus Modeller.

In **Figure 5**, we presented four representative targets in the AIDA dataset that SAXSDom generated accurate models compared to the modeling methods without using SAXS information. In **Figure 5(A)**, the crystal structure of *signal recognition particle receptor from E.coli* - 1ftsA (chain A of 1FTS) contains one *α*-helix domain (residues 1-82) linked by another *α*-helix+*β*-sheet+*α*-helix domain (residues 92-295) with linker length consisting of 9 residues. SAXSDom successfully folded the domains into correct orientation using SAXS information and the assembled domains were largely overlaid with the envelope of the protein structure even though the variation of linker region is relatively large. The predicted assembly model has RMSD= 2.776, TMscore=0.876 against the native structure, and *χ* score=2.774 with matching to the SAXS profile. **Figure 6** shows the comparison of predicted models by SAXSDom and three modeling methods. **Figure 6(A)** shows that the correct orientation of domains predicted by SAXSDom results in a better matching to the SAXS data than other methods in the both pair distance distribution and scattering curves. **Figure 6(B)** shows that the first *α*-helix domain was oriented to correct position in the full-length structure with slight bias. The residue-specific distance errors between the native structure and the models predicted by four methods are shown in **Figure 6(C)**. The case shows that the accuracy of domain assembly was improved by incorporating SAXS energies in the SAXSDom compared to *ab initio* method SAXSDom-abinitio.

**Figure 5.**
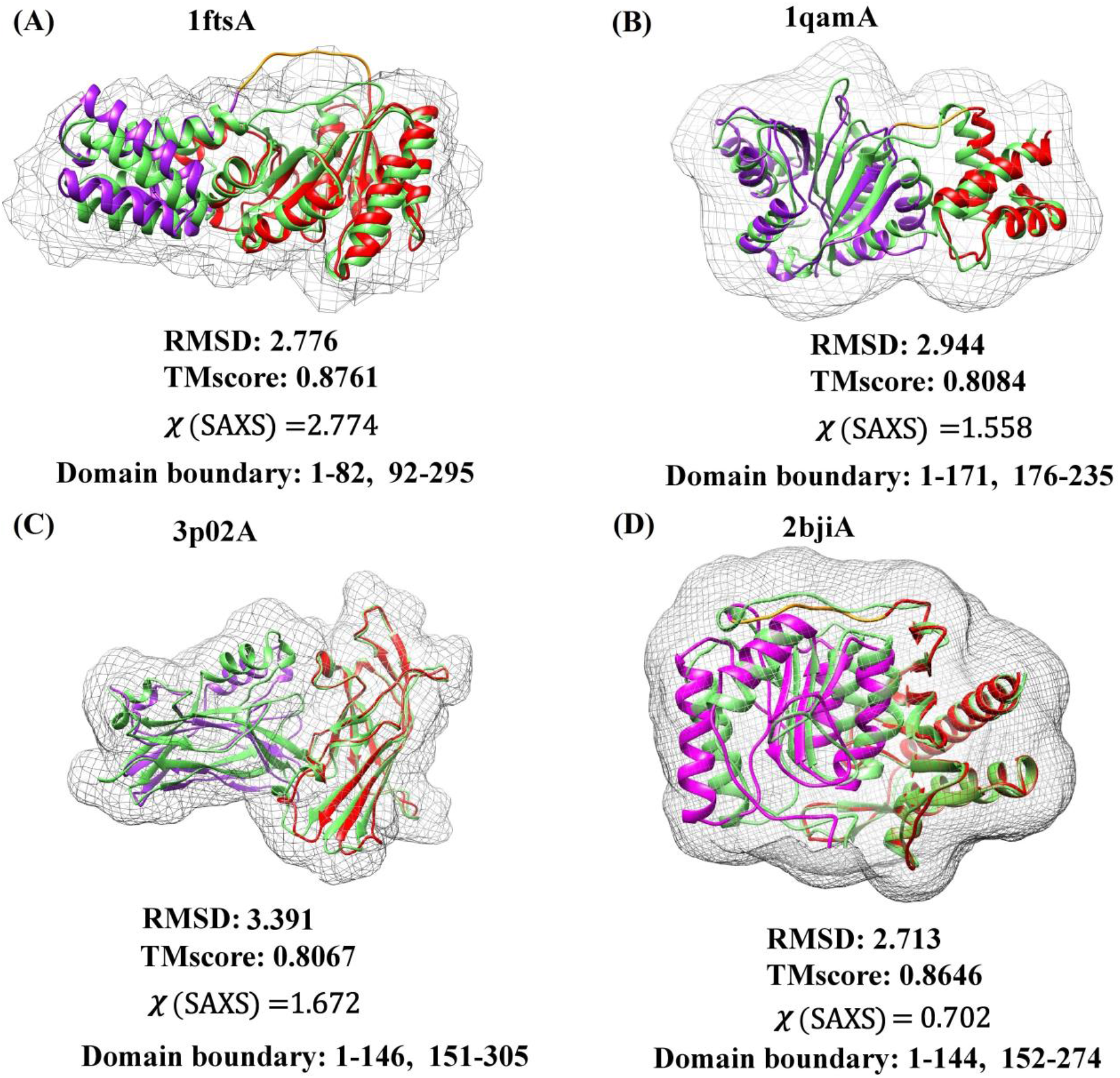
The predicted assembly models and shape envelops of five two-domain proteins. The predicted model (colored) and the native structure (green) is superimposed. The domain linker (yellow) and domains (purple, red) are highlighted in the predicted model. (A) The signal recognition particle receptor from E.coli - 1ftsA (chain A of 1FTS), linker length = 9, RMSD=2.776, TMscore=0.876, *χ* score=2.774. (B) A structure of the rRNA methyltransferase ErmC’ - 1qamA (chain A of 1QAM), linker length = 4, RMSD=2.944, TMscore=0.808, *χ* score=1.558. (C) A structure of from Bacteroides ovatus – 3p02A (chain A of 3P02), linker length = 4, RMSD=3.391, TMscore=0.8067, *χ* score=1.672. (D) Myo-inositol monophosphatase enzyme – 2bji (chain A of 2BJI), linker length = 7, RMSD=2.713, TMscore=0.8646, *χ* score=0.702.

**Figure 6.**
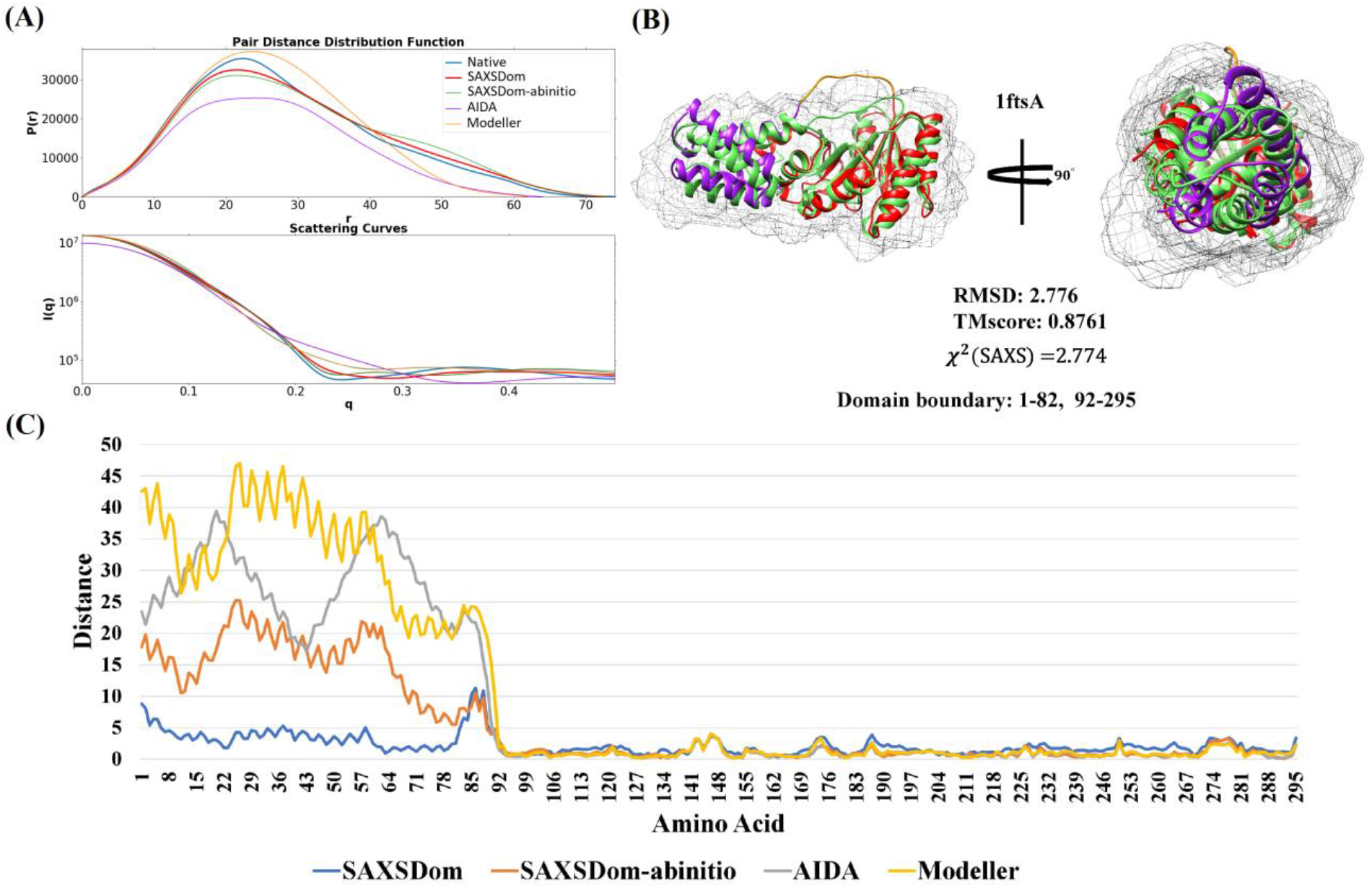
Comparison of predicted models for protein 1ftsA (chain A of 1FTS) by SAXSDom, AIDA and Modeller. (A) Fittings of SAXS profiles between the theoretical SAXS data and computed SAXS data of predicted models. The fitting curves of native data (blue) and SAXSDom model (red) are highlighted as thick lines while the curves of other three methods are represented as thick lines. (B) The assembled full-length model with quality measurements. (C) Residue-specific distance error between the predicted models and the native structure.

**Figure 5(B)** shows the predicted domain assembly for ErmC’ protein (PDB entry 1QAM). The structure consists of two domains, an N-terminal *α*-helix+*β*-sheet+*α*-helix domain (residues 1-171) and a C-terminal *α*-helix domain (residues 176-235). The predicted assembly model has RMSD= 2.994, TMscore=0.8084 to the native structure, and *χ* score=1.558 to the SAXS profile. The domain linker contains 4 residues and is folded into similar shape as that in the native structure. Domain assembly for protein 3po2A also achieved good performance, as shown in **Figure 5(C)**, and the domains are folded to native-like orientation with RMSD=3.391, TMscore=0.8067 and *χ* score=1.672.

**Figure 5(D)** presents the predicted assembly for protein 2bjiA. The full-length structure of 2bjiA is folded as a penda-layered *α*-helix+*β*-sheet+*α*-helix+*β*-sheet+*α*-helix sandwich, and the linker connects the first N-terminal *α*-helix+*β*-sheet domain (residues 1-144) with second C-terminal - helix+*β*-sheet+α-helix domain (residues 152-257). SAXSDom successfully generated a well-folded model that domains were oriented to a native-like state with RMSD=2.713, TMscore=0.8646 and *χ* score=0.702. The detailed results of different domain assembly methods for proteins qamA, 3po2A, and 2bjiA are visualized in **Figure S5, S6 and S7**.

## 4. Conclusion and Future work

In this work, we developed a data-assisted *ab initio* domain assembly method, SAXSDom, by integrating the probabilistic approach for backbone conformation sampling with SAXS-assisted restraints in domain assembly. Our method designed and evaluated several SAXS-related score functions for structure modeling by extracting useful restraints from theoretical SAXS data in different aspects, including fitness of SAXS intensities, the divergence of pair-atom distance distribution, and agreement of radius of gyration derived from the SAXS data and the model. Our results show that incorporating the restraints from SAXS data into de novo conformational sampling method can improve the protein domain assembly. SAXSDom generates more accurate domain assembly for 40 out of 46 CASP multi-domain proteins, when models generated with and without SAXS information are evaluated using RMSD and TMscore. On the AIDA dataset, SAXSDom achieved higher accuracy for at least 45 out of 73 multi-domain proteins. We believe that our method can be further improved in several ways: (1) adopting new physical energies derived from full-atom structures such as van der Waals hard sphere repulsion, residue environment, residue pair, radius of gyration as introduced in Rosetta ^11^; (2) extending the continuous domain assembly with discontinuous domain assembly for those proteins with inserted domains; (3) designing more advanced SAXS scoring functions to guide the structure folding.

## 5. Acknowledgements

This work is supported by an NIH grant (R01GM093123), an NSF IIS grant (IIS1763246), and an NSF DBI grant (DBI1759934) to JC.

## Reference

1. Rost, B. Did evolution leap to create the protein universe? Current opinion in structural biology 2002;12(3):409–416.

2. Jaenicke, R. Folding and association of proteins. Progress in biophysics and molecular biology 1987;49(2–3): 117–237.

3. McLachlan, A. Gene duplications in the structural evolution of chymotrypsin. Journal of molecular biology 1979;128(1):49–79.

4. Berrondo, M.; Ostermeier, M.; Gray, J. J. Structure prediction of domain insertion proteins from structures of individual domains. Structure 2008;16(4):513–527.

5. Robinson, C. R.; Sauer, R. T. Optimizing the stability of single-chain proteins by linker length and composition mutagenesis. Proceedings of the National Academy of Sciences 1998;95(11):5929–5934.

6. George, R. A.; Heringa, J. An analysis of protein domain linkers: their classification and role in protein folding. Protein Engineering, Design and Selection 2002;15(11):871–879.

7. Dueber, J. E.; Yeh, B. J.; Chak, K.; Lim, W. A. Reprogramming control of an allosteric signaling switch through modular recombination. Science 2003;301(5641):1904–1908.

8. Kim, D. E.; Chivian, D.; Baker, D. Protein structure prediction and analysis using the Robetta server. Nucleic acids research 2004;32(suppl_2):W526–W531.

9. Li, J.; Adhikari, B.; Cheng, J. An improved integration of template-based and template-free protein structure modeling methods and its assessment in CASP11. Protein and peptide letters 2015;22(7):586–593.

10. Rose, P. W.; Prlić, A.; Bi, C.; Bluhm, W. F.; Christie, C. H.; Dutta, S.; Green, R. K.; Goodsell, D. S.; Westbrook, J. D.; Woo, J. The RCSB Protein Data Bank: views of structural biology for basic and applied research and education. Nucleic acids research 2014;43(D1):D345–D356.

11. Rohl, C. A.; Strauss, C. E.; Misura, K. M.; Baker, D. Protein structure prediction using Rosetta. Methods in enzymology 2004;383:66–93.

12. Yang, J.; Yan, R.; Roy, A.; Xu, D.; Poisson, J.; Zhang, Y. The I-TASSER Suite: protein structure and function prediction. Nature methods 2015;12(1):7.

13. Adhikari, B.; Bhattacharya, D.; Cao, R.; Cheng, J. CONFOLD: residue-residue contact-guided ab initio protein folding. Proteins: Structure, Function, and Bioinformatics 2015;83(8): 1436–1449.

14. Adhikari, B.; Cheng, J. CONFOLD2: Improved contact-driven ab initio protein structure modeling. bioRxiv 2017:228460.

15. Hou, J.; Adhikari, B.; Cheng, J. DeepSF: deep convolutional neural network for mapping protein sequences to folds. arXiv preprint arXiv:1706.01010 2017.

16. Källberg, M.; Margaryan, G.; Wang, S.; Ma, J.; Xu, J., RaptorX server: a resource for template-based protein structure modeling. In Protein Structure Prediction, Springer: 2014; pp 17–27.

17. Jo, T.; Hou, J.; Eickholt, J.; Cheng, J. Improving protein fold recognition by deep learning networks. Scientific reports 2015;5:srep17573.

18. Krieger, E.; Nabuurs, S. B.; Vriend, G. Homology modeling. Methods of biochemical analysis 2003;44:509–524.

19. Bhattacharya, D.; Cao, R.; Cheng, J. UniCon3D: de novo protein structure prediction using united-residue conformational search via stepwise, probabilistic sampling. Bioinformatics 2016;32(18):2791–2799.

20. Eswar, N.; Webb, B.; Marti-Renom, M. A.; Madhusudhan, M.; Eramian, D.; Shen, M. y.; Pieper, U.; Sali, A. Comparative protein structure modeling using Modeller. Current protocols in bioinformatics 2006;15(1):5.6. 1–5.6. 30.

21. Xu, D.; Jaroszewski, L.; Li, Z.; Godzik, A. AIDA: ab initio domain assembly for automated multi-domain protein structure prediction and domain–domain interaction prediction. Bioinformatics 2015;31(13):2098–2105.

22. Belsom, A.; Schneider, M.; Brock, O.; Rappsilber, J. Blind evaluation of hybrid protein structure analysis methods based on cross-linking. Trends in biochemical sciences 2016;41(7):564–567.

23. Ogorzalek, T. L.; Hura, G. L.; Belsom, A.; Burnett, K. H.; Kryshtafovych, A.; Tainer, J. A.; Rappsilber, J.; Tsutakawa, S. E.; Fidelis, K. Small angle X-ray scattering and cross-linking for data assisted protein structure prediction in CASP 12 with prospects for improved accuracy. Proteins: Structure, Function, and Bioinformatics 2018;86:202–214.

24. Moult, J.; Fidelis, K.; Kryshtafovych, A.; Schwede, T.; Tramontano, A. Critical assessment of methods of protein structure prediction (CASP)—Round XII. Proteins: Structure, Function, and Bioinformatics 2018;86:7–15.

25. Korasick, D. A.; Tanner, J. J. Determination of protein oligomeric structure from small-angle X-ray scattering. Protein Science 2018;27(4):814–824.

26. Li, J.; Jiao, A.; Chen, S.; Wu, Z.; Xu, E.; Jin, Z. Application of the small-angle X-ray scattering technique for structural analysis studies: A review. Journal of Molecular Structure 2018;1165:391–400.

27. Wriggers, W. Using Situs for the integration of multi-resolution structures. Biophysical reviews 2010;2(1):21–27.

28. Schneidman-Duhovny, D.; Hammel, M.; Tainer, J. A.; Sali, A. Accurate SAXS profile computation and its assessment by contrast variation experiments. Biophysical journal 2013;105(4):962–974.

29. Franke, D.; Petoukhov, M.; Konarev, P.; Panjkovich, A.; Tuukkanen, A.; Mertens, H.; Kikhney, A.; Hajizadeh, N.; Franklin, J.; Jeffries, C. ATSAS 2.8: a comprehensive data analysis suite for small-angle scattering from macromolecular solutions. Journal of applied crystallography 2017;50(4):1212–1225.

30. Dos Reis, M. A.; Aparicio, R.; Zhang, Y. Improving protein template recognition by using small-angle x-ray scattering profiles. Biophysical journal 2011;707(11):2770–2781.

31. Joo, K.; Heo, S.; Joung, I.; Hong, S. H.; Lee, S. J.; Lee, J. Data-assisted protein structure modeling by global optimization in CASP12. Proteins: Structure, Function, and Bioinformatics 2018;86:240–246.

32. Ogorzalek, T. L.; Hura, G. L.; Kryshtafovych, A.; Tainer, J. A.; Fidelis, K.; Tsutakawa, S. E. Small Angle X-ray Scattering for Data-Assisted Structure Prediction in CASP12 with Prospects to Improve Accuracy. Biophysical Journal 2018;774(3):576a–577a.

33. Jiménez-García, B.; Pons, C.; Svergun, D. I.; Bernadó, P.; Fernández-Recio, J. pyDockSAXS: protein-protein complex structure by SAXS and computational docking. Nucleic acids research 2015;43(W1):W356–W361.

34. Zhang, Y.; Skolnick, J. Scoring function for automated assessment of protein structure template quality. Proteins: Structure, Function, and Bioinformatics 2004;57(4):702–710.

35. Kryshtafovych, A.; Monastyrskyy, B.; Fidelis, K. CASP 11 statistics and the prediction center evaluation system. Proteins: Structure, Function, and Bioinformatics 2016;84:15–19.

36. Tress, M. L.; Ezkurdia, I.; Richardson, J. S. Target domain definition and classification in CASP8. Proteins: Structure, Function, and Bioinformatics 2009;77(S9):10–17.

37. Xu, Y.; Xu, D.; Gabow, H. N. Protein domain decomposition using a graph-theoretic approach. Bioinformatics 2000;16(12): 1091–1104.

38. Hou, J.; Wu, T.; Cao, R.; Cheng, J. Protein tertiary structure modeling driven by deep learning and contact distance prediction in CASP13. bioRxiv 2019:552422.

39. Liwo, A.; Ołdziej, S.; Pincus, M. R.; Wawak, R. J.; Rackovsky, S.; Scheraga, H. A. A united-residue force field for off-lattice protein-structure simulations. I. Functional forms and parameters of long-range side-chain interaction potentials from protein crystal data. Journal of computational chemistry 1997;18(7):849–873.

40. Magnan, C. N.; Baldi, P. SSpro/ACCpro 5: almost perfect prediction of protein secondary structure and relative solvent accessibility using profiles, machine learning and structural similarity. Bioinformatics 2014;30(18):2592–2597.

41. Dill, K. A. Dominant forces in protein folding. Biochemistry 1990;29(31):7133–7155.

42. Schneidman-Duhovny, D.; Hammel, M.; Sali, A. FoXS: a web server for rapid computation and fitting of SAXS profiles. Nucleic acids research 2010;38(suppl_2):W540–W544.

43. Liu, H.; Zwart, P. H. Determining pair distance distribution function from SAXS data using parametric functionals. Journal of structural biology 2012;180(1):226–234.

44. Russel, D.; Lasker, K.; Webb, B.; Velázquez-Muriel, J.; Tjioe, E.; Schneidman-Duhovny, D.; Peterson, B.; Sali, A. Putting the pieces together: integrative modeling platform software for structure determination of macromolecular assemblies. PLoS biology 2012;10(1):e1001244.

45. Rotkiewicz, P.; Skolnick, J. Fast procedure for reconstruction of full-atom protein models from reduced representations. Journal of computational chemistry 2008;29(9): 1460–1465.

46. Koehler Leman, J.; Bonneau, R. A novel domain assembly routine for creating full-length models of membrane proteins from known domain structures. Biochemistry 2017;57(13): 1939–1944.

47. Svergun, D. Determination of the regularization parameter in indirect-transform methods using perceptual criteria. Journal of applied crystallography 1992;25(4):495–503.

48. Cheng, J.; Randall, A. Z.; Sweredoski, M. J.; Baldi, P. SCRATCH: a protein structure and structural feature prediction server. Nucleic acids research 2005;33(suppl_2):W72–W76.

49. Cao, R.; Cheng, J. Protein single-model quality assessment by feature-based probability density functions. Scientific reports 2016;6:23990.

