## Supplemental File for "SAXSDom: Modeling multi-domain protein structures using small-angle X-ray scattering data"

#### Tables

| **CASP** | **# of Proteins** | **# of Models** |
|---|---|---|
| CASP8 | 123 | 35689 |
| CASP9 | 117 | 34998 |
| CASP10 | 103 | 25349 |
| CASP11 | 85 | 16014 |
| Total | 428 | 112050 |

**Table S1**. Statistics of the collected proteins and predicted server models from CASP8-11.

| Scores | Score Function | Dataset | | | | |
|---|---|---|---|---|---|---|
| | | CASP8 | CASP9 | CASP10 | CASP11 | Average |
| **1** | $\sqrt{\frac{1}{N}\sum_{i=1}^{N}(\frac{I_{exp}(q_i) - cI_{model}(q_i)}{\sigma(q_i)})^2}$ | **0.25** | **0.49** | **0.54** | **0.60** | **0.47** |

| | | | | | | |
|---|---|---|---|---|---|---|
| **2** | $\boldsymbol{\sum_{i=1}^{N} Pr_{model}(r_i) * log \frac{Pr_{model}(r_i)}{Pr_{exp}(r_i)}}$ | **0.48** | **0.69** | **0.58** | **0.65** | **0.60** |
| **3** | $\boldsymbol{\frac{\lvert RG_{exp} - RG_{model}\rvert}{\lvert RG_{exp}\rvert}}$ | **0.54** | **0.81** | **0.69** | **0.75** | **0.70** |
| 4 | $\frac{1}{N}\sum_{i=1}^{N}(\frac{I_{exp}(q_i) - I_{model}(q_i)}{\sigma(q_i)})^2$ | 0.13 | 0.37 | 0.48 | 0.53 | 0.38 |
| **5** | $\boldsymbol{\frac{\sum_{i=1}^{N}\lvert I_{exp}(q_i) - I_{model}(q_i)\rvert}{\sum_{i=1}^{N}\lvert I_{exp}(q_i)\rvert}}$ | **0.43** | **0.65** | **0.61** | **0.67** | **0.59** |
| 6 | $\frac{\sum_{i=1}^{N}\lvert \log(I_{exp}(q_i)) - \log(I_{model}(q_i))\rvert}{\sum_{i=1}^{N}\lvert \log(I_{exp}(q_i))\rvert}$ | 0.30 | 0.41 | 0.42 | 0.46 | 0.40 |
| 7 | $\sum_{i=1}^{N} q_i * (I_{exp}(q_i) - I_{model}(q_i))^2$ | 0.32 | 0.59 | 0.56 | 0.61 | 0.52 |
| 8 | $\sum_{i=1}^{N} q_i * (\log(I_{exp}(q_i)) - \log(I_{model}(q_i)))^2$ | 0.18 | 0.24 | 0.29 | 0.32 | 0.25 |
| 9 | $\frac{\sum_{i=1}^{N} {q_i}^2\lvert I_{exp}(q_i) - I_{model}(q_i)\rvert}{\sum_{i=1}^{N} {q_i}^2\lvert I_{exp}(q_i)\rvert}$ | 0.25 | 0.35 | 0.41 | 0.49 | 0.37 |
| 10 | $\sum_{i=1}^{N}(Pr_{exp}(r_i) - Pr_{model}(r_i))^2$ | 0.26 | 0.48 | 0.40 | 0.52 | 0.41 |
| 11 | $log\ (1 - corr(I_{exp}(q_i), I_{model}(q_i)))$ | 0.46 | 0.56 | 0.51 | 0.55 | 0.52 |
| 12 | $log\ (1 - corr(Pr_{exp}(r_i), Pr_{model}(r_i)))$ | 0.41 | 0.49 | 0.43 | 0.49 | 0.46 |
| 13 | $log\ (1 - cos(I_{exp}(q_i), I_{model}(q_i)))$ | 0.46 | 0.56 | 0.51 | 0.55 | 0.52 |
| 14 | $log\ (1 - cos(Pr_{exp}(r_i), Pr_{model}(r_i)))$ | 0.42 | 0.50 | 0.44 | 0.49 | 0.46 |

**Table S2**. Pearson's correlation of the structural quality (RMSD) against the SAXS score functions derived from Full-atom of protein structure. Analysis was done based on the predicted models from CASP8-11

| Score | Score Function | Dataset | | | | |
|---|---|---|---|---|---|---|
| | | CASP8 | CASP9 | CASP10 | CASP11 | Average |
| **1** | $\boldsymbol{\sqrt{\frac{1}{N}\sum_{i=1}^{N}(\frac{I_{exp}(q_i) - cI_{model}(q_i)}{\sigma(q_i)})^2}}$ | **0.20** | **0.42** | **0.42** | **0.49** | **0.38** |
| **2** | $\boldsymbol{\sum_{i=1}^{N} Pr_{model}(r_i) * log \frac{Pr_{model}(r_i)}{Pr_{exp}(r_i)}}$ | **0.45** | **0.68** | **0.56** | **0.63** | **0.58** |
| **3** | $\boldsymbol{\frac{\lvert RG_{exp} - RG_{model}\rvert}{\lvert RG_{exp}\rvert}}$ | **0.47** | **0.80** | **0.64** | **0.72** | **0.66** |

| | | | | | | |
|---|---|---|---|---|---|---|
| 4 | $\frac{1}{N}\sum_{i=1}^{N}(\frac{I_{exp}(q_i)-I_{model}(q_i)}{\sigma(q_i)})^2$ | 0.10 | 0.34 | 0.36 | 0.44 | 0.31 |
| **5** | $\boldsymbol{\frac{\sum_{i=1}^{N}\lvert I_{exp}(q_i)-I_{model}(q_i)\rvert}{\sum_{i=1}^{N}\lvert I_{exp}(q_i)\rvert}}$ | **0.33** | **0.59** | **0.54** | **0.61** | **0.52** |
| 6 | $\frac{\sum_{i=1}^{N}\lvert\log(I_{exp}(q_i))-\log(I_{model}(q_i))\rvert}{\sum_{i=1}^{N}\lvert\log(I_{exp}(q_i))\rvert}$ | 0.25 | 0.38 | 0.36 | 0.42 | 0.35 |
| 7 | $\sum_{i=1}^{N}q_i*(I_{exp}(q_i)-I_{model}(q_i))^2$ | 0.24 | 0.55 | 0.50 | 0.58 | 0.47 |
| 8 | $\sum_{i=1}^{N}q_i*(\log(I_{exp}(q_i))-\log(I_{model}(q_i)))^2$ | 0.15 | 0.24 | 0.26 | 0.33 | 0.25 |
| 9 | $\frac{\sum_{i=1}^{N}{q_i}^2\lvert I_{exp}(q_i)-I_{model}(q_i)\rvert}{\sum_{i=1}^{N}{q_i}^2\lvert I_{exp}(q_i)\rvert}$ | 0.16 | 0.27 | 0.28 | 0.36 | 0.27 |
| 10 | $\sum_{i=1}^{N}(Pr_{exp}(r_i)-Pr_{model}(r_i))^2$ | 0.23 | 0.47 | 0.36 | 0.50 | 0.39 |
| 11 | $log\ (1-\ corr(I_{exp}(q_i),I_{model}(q_i)))$ | 0.38 | 0.52 | 0.45 | 0.49 | 0.46 |
| 12 | $log\ (1-corr(Pr_{exp}(r_i),Pr_{model}(r_i)))$ | 0.36 | 0.46 | 0.40 | 0.46 | 0.42 |
| 13 | $log\ (1-\ cos(I_{exp}(q_i),I_{model}(q_i)))$ | 0.37 | 0.52 | 0.45 | 0.50 | 0.46 |
| 14 | $log\ (1-cos(Pr_{exp}(r_i),Pr_{model}(r_i)))$ | 0.37 | 0.47 | 0.41 | 0.46 | 0.43 |

**Table S3**. Pearson's correlation of the structural quality (RMSD) against the SAXS score functions derived from Ca Atom of protein structure. Analysis was done based on the predicted models from CASP8-11.

| **ScoreFunction** | **Top 1** | | **Best-of-five** | | **Best-of-50** | |
|---|---|---|---|---|---|---|
| | **TM-score** | **RMSD** | **TM-score** | **RMSD** | **TM-score** | **RMSD** |
| SAXSDom-abinitio | | | | | | |
| SAXSDom ($E_{saxs\cdot\chi}$) | 2.86E-05 | 1.32E-04 | 6.53E-06 | 7.02E-05 | 2.09E-05 | 6.21E-04 |
| SAXSDom ($E_{saxs\cdot IntFit}$) | 3.98E-02 | 3.84E-02 | 3.69E-04 | 5.45E-04 | 9.67E-05 | 1.66E-03 |
| SAXSDom ($E_{saxs\cdot Pr}$) | 2.82E-04 | 7.67E-04 | 4.57E-05 | 1.25E-04 | 7.01E-06 | 1.29E-04 |
| SAXSDom ($E_{saxs\cdot Rg}$) | 5.99E-03 | 5.10E-03 | 4.32E-03 | 3.41E-03 | 3.42E-03 | 7.60E-03 |
| SAXSDom ($E_{saxs}$) | 2.38E-04 | 3.21E-04 | 3.08E-05 | 1.39E-04 | 5.62E-07 | 7.84E-05 |

**Table S4**. P-value of one-tailed paired t-test of TM-score and RMSD between different SAXS-related score functions and ab-initio potential.

### Figures

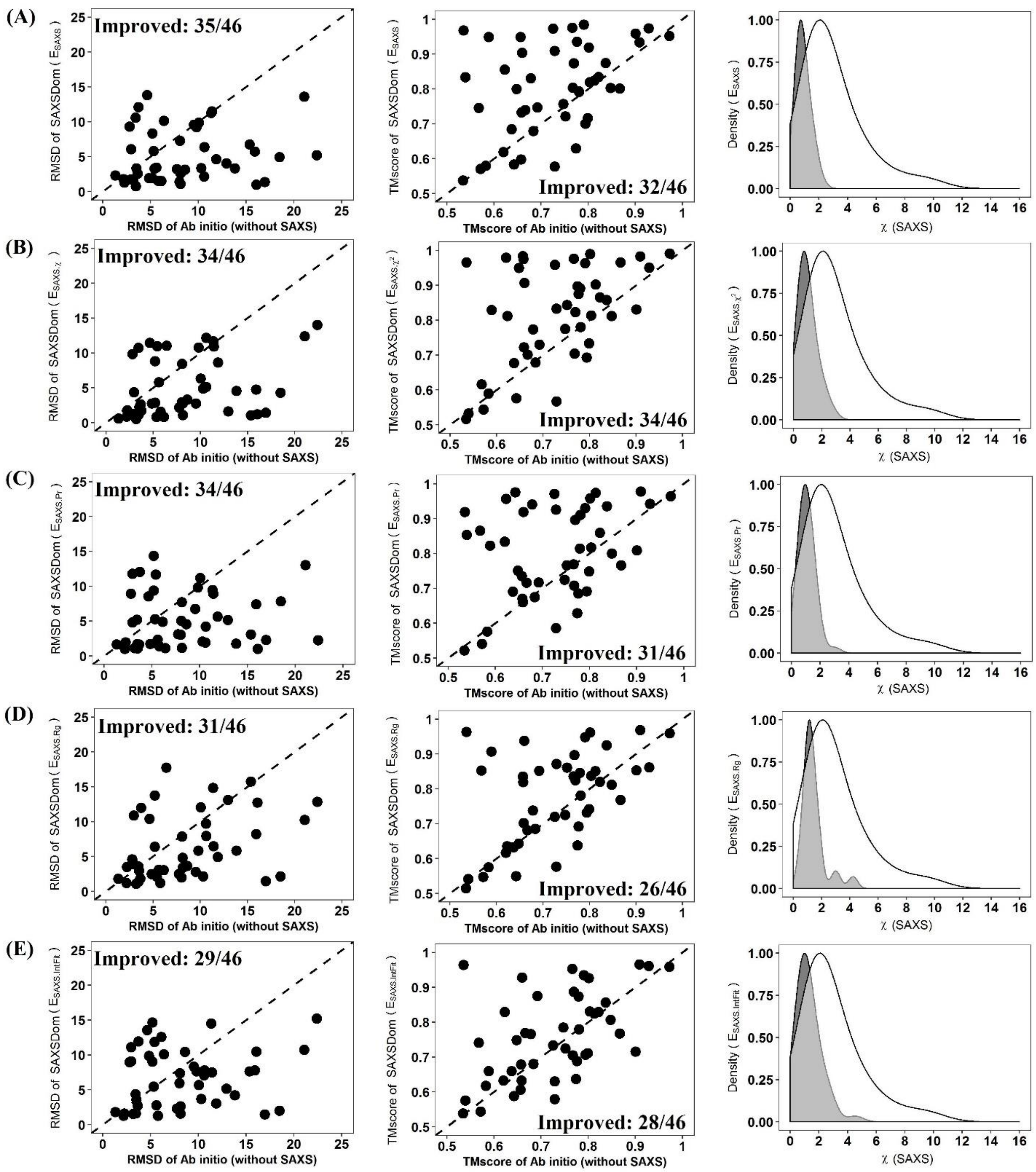


**Figure S1**. Comparison of five SAXSDom methods with SAXSDom-abinitio method on the top one assembled model. **(A)** SAXSDom ($E_{saxs}$) versus SAXSDom-abinitio (Left plot: TM_scores of SAXSDom ($E_{saxs}$), models versus TM_scores of SAXSDom-abinitio models; Middle plot: RMSD of the models of the two methods; Right plot: Distribution of χ score of all assembled models for 46 proteins by two methods). **(B)** SAXSDom ($E_{saxs\cdot\chi}$) versus SAXSDom-abinitio. **(C)** SAXSDom ($E_{saxs\cdot Pr}$) versus SAXSDom-abinitio. **(D)** SAXSDom ($E_{saxs\cdot Rg}$) versus SAXSDom-abinitio. **(E)** SAXSDom ($E_{saxs\cdot IntFit}$) versus SAXSDom-abinitio.

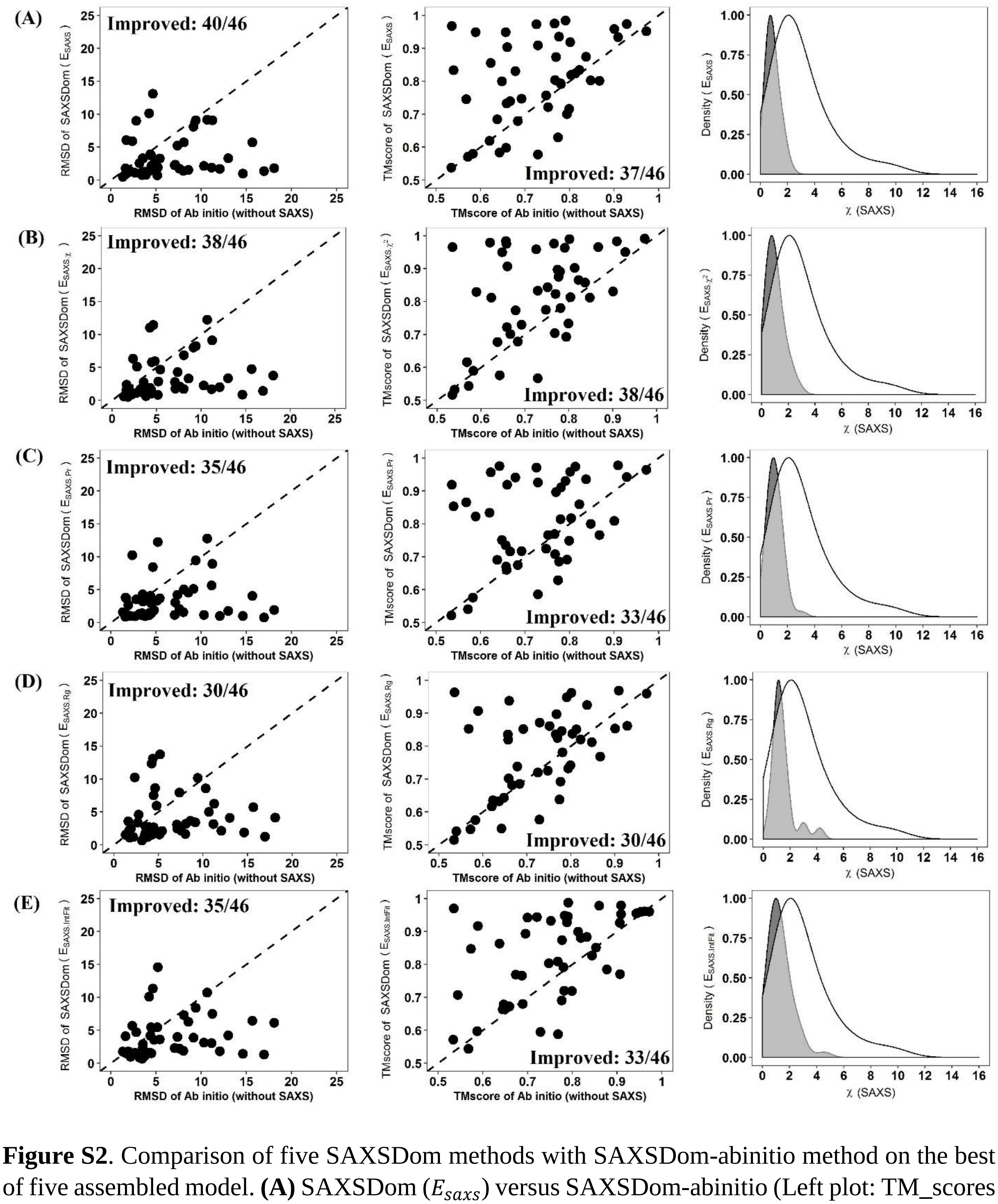


**Figure S2**. Comparison of five SAXSDom methods with SAXSDom-abinitio method on the best of five assembled model. **(A)** SAXSDom ($E_{saxs}$) versus SAXSDom-abinitio (Left plot: TM_scores of SAXSDom ($E_{saxs}$), models versus TM_scores of SAXSDom-abinitio models; Middle plot: RMSD of the models of the two methods; Right plot: Distribution of $\chi$ score of all assembled models for 46 proteins by two methods). **(B)** SAXSDom ($E_{saxs\cdot\chi}$) versus SAXSDom-abinitio. **(C)**

SAXSDom ($E_{saxs \cdot Pr}$) versus SAXSDom-abinitio. **(D)** SAXSDom ($E_{saxs \cdot Rg}$) versus SAXSDom-abinitio. **(E)** SAXSDom ($E_{saxs \cdot IntFit}$) versus SAXSDom-abinitio.

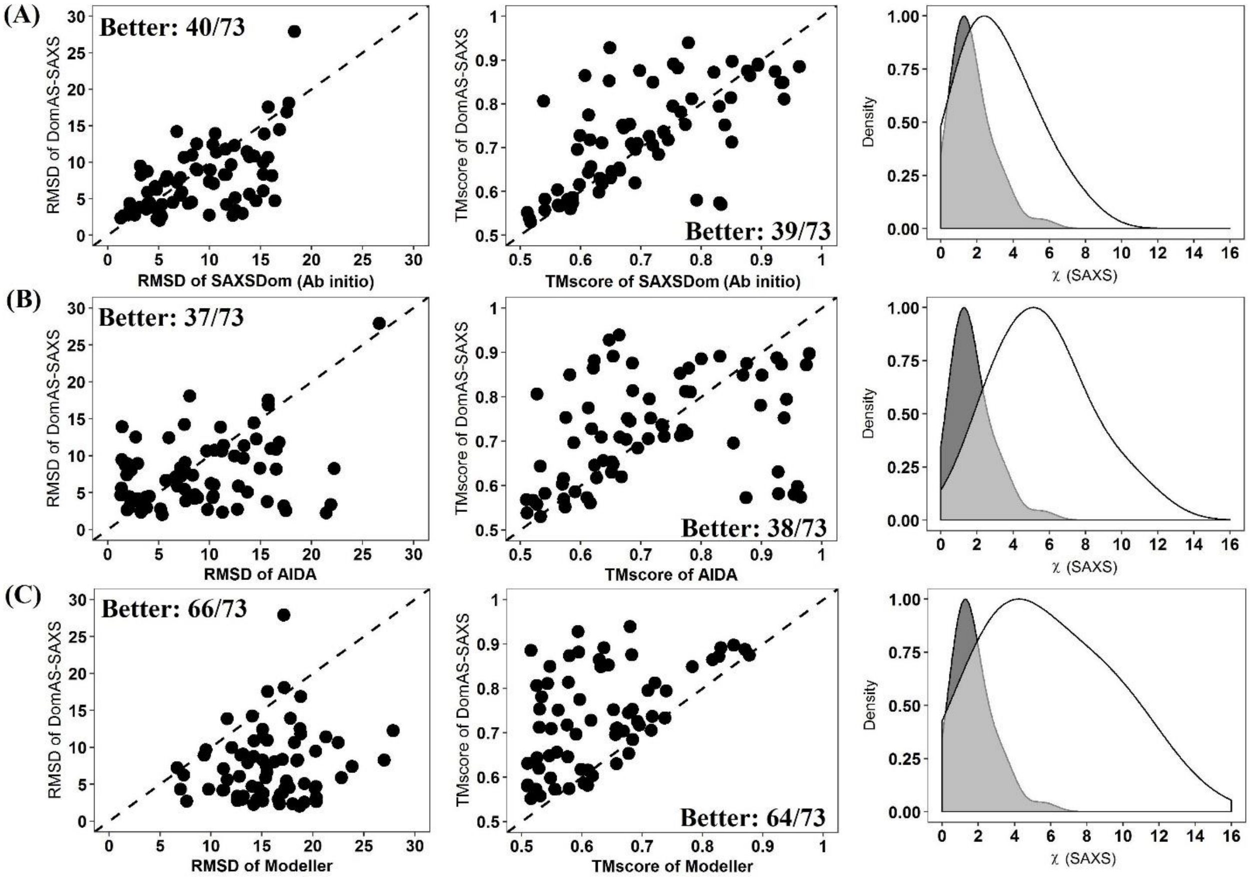


**Figure S3**. Comparison of five SAXSDom methods with SAXSDom-abinitio, AIDA and Modeller on the top 1 assembled model. **(A)** SAXSDom versus SAXSDom-abinitio (Left plot: TM_scores of SAXSDom, models versus TM_scores of SAXSDom-abinitio models; Middle plot: RMSD of the models of the two methods; Right plot: Distribution of χ score of all assembled models for 46 proteins by two methods). **(B)** SAXSDom versus AIDA. **(C)** SAXSDom versus Modeller.

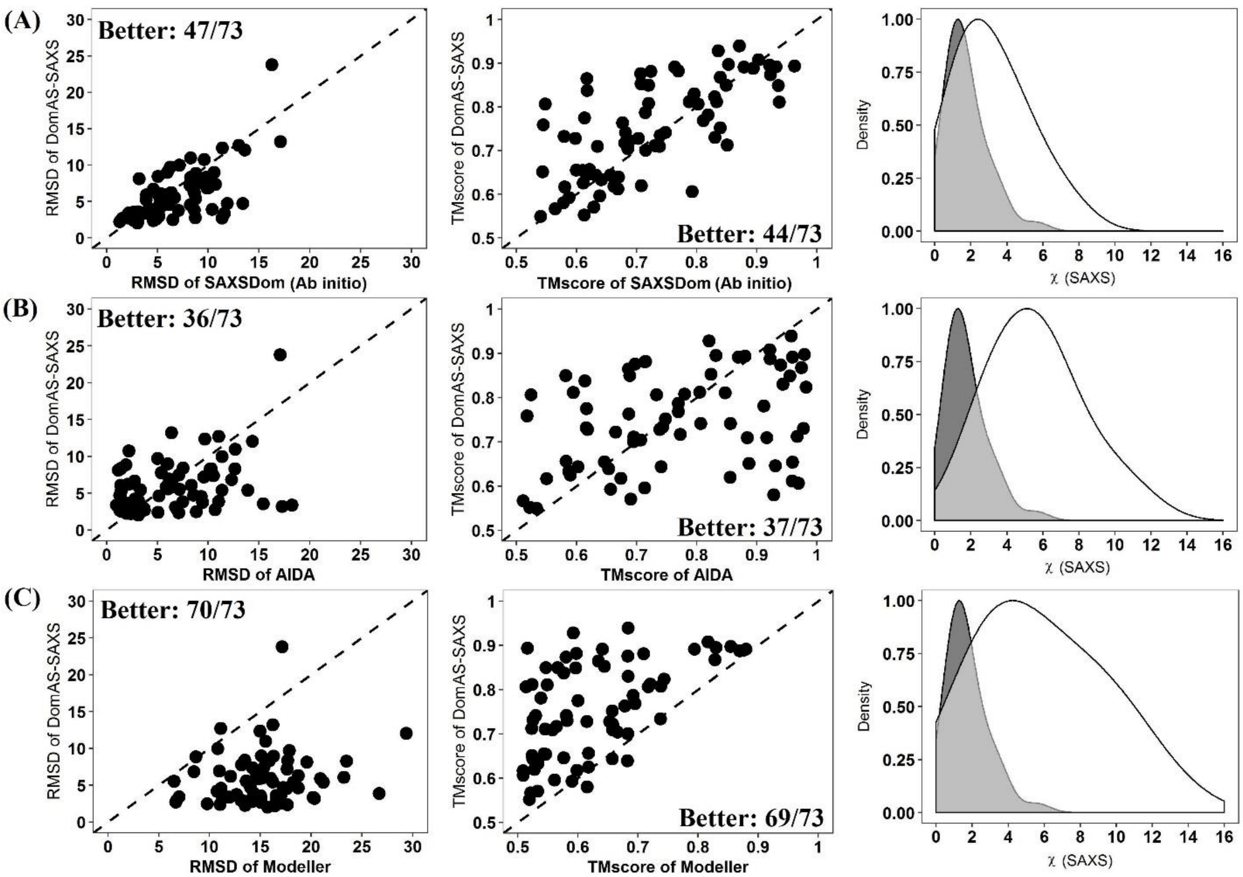


**Figure S4**. Comparison of five SAXSDom methods with SAXSDom-abinitio, AIDA and Modeller on the best of 5 assembled model. **(A)** SAXSDom versus SAXSDom-abinitio (Left plot: TM_scores of SAXSDom, models versus TM_scores of SAXSDom-abinitio models; Middle plot: RMSD of the models of the two methods; Right plot: Distribution of χ score of all assembled models for 46 proteins by two methods). **(B)** SAXSDom versus AIDA. **(C)** SAXSDom versus Modeller.

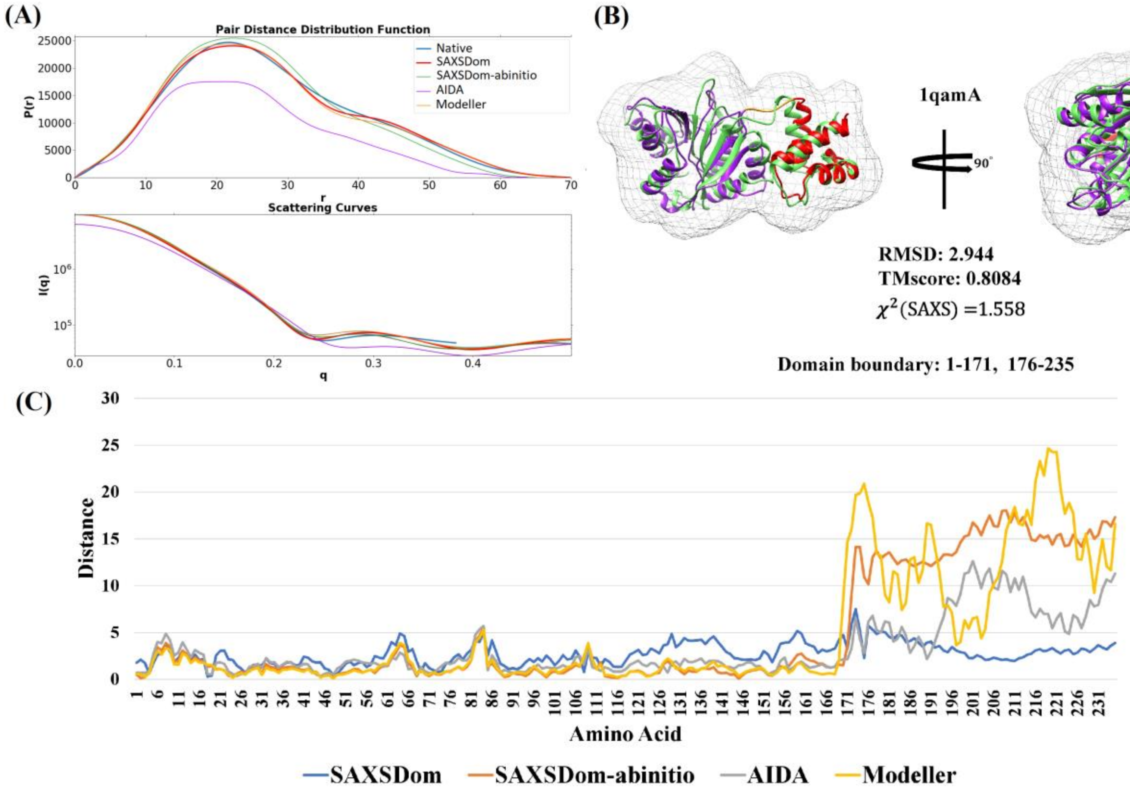


**Figure S5.** Comparison of predicted models for protein 1qamA (chain A of 1QAM) by SAXSDom, AIDA and Modeller. (A) Fittings of SAXS profiles between the theoretical SAXS data and computed SAXS data of predicted models. The fitting curves of native data (blue) and SAXSDom model (red) are highlighted as thick lines while the curves of other three methods are represented as thick lines. (B) The assembled full-length model with quality measurements. (C) Residue-specific distance error between the predicted models and the native structure.

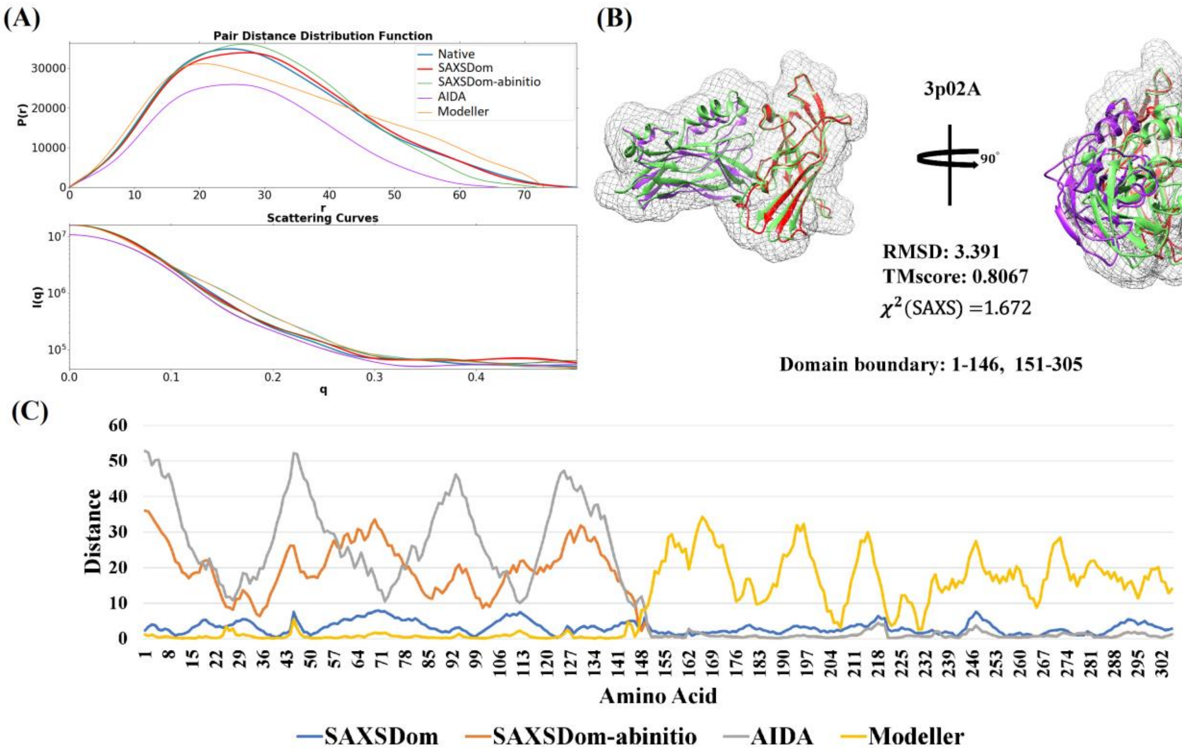


**Figure S6.** Comparison of predicted models for protein 3p02A (chain A of 3P02) by SAXSDom, AIDA and Modeller. (A) Fittings of SAXS profiles between the theoretical SAXS data and computed SAXS data of predicted models. The fitting curves of native data (blue) and SAXSDom model (red) are highlighted as thick lines while the curves of other three methods are represented as thick lines. (B) The assembled full-length model with quality measurements. (C) Residue-specific distance error between the predicted models and the native structure.

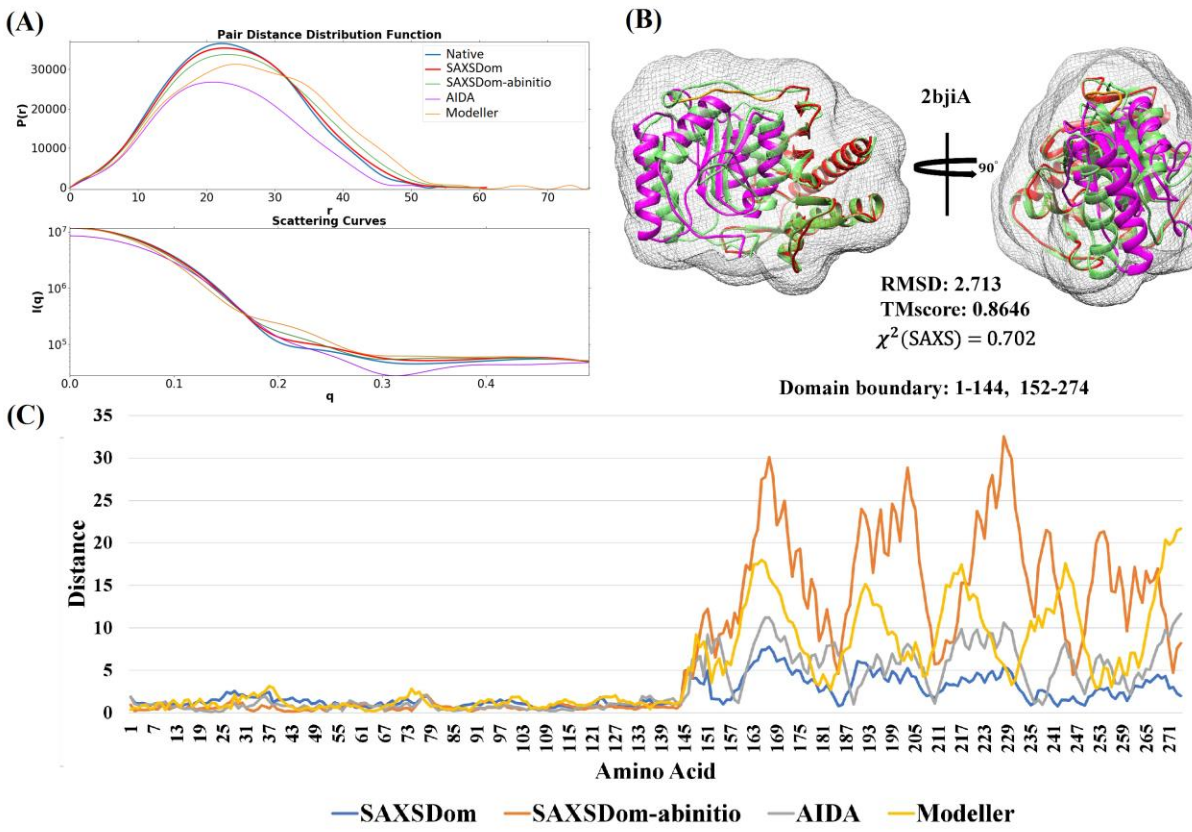


**Figure S7.** Comparison of predicted models for protein 2bjiA (chain A of 2BJI) by SAXSDom, AIDA and Modeller. (A) Fittings of SAXS profiles between the theoretical SAXS data and computed SAXS data of predicted models. The fitting curves of native data (blue) and SAXSDom model (red) are highlighted as thick lines while the curves of other three methods are represented as thick lines. (B) The assembled full-length model with quality measurements. (C) Residue-specific distance error between the predicted models and the native structure.
